# Nuclear Expression and DNA Binding Capacity of Receptor for Advanced Glycation End Products in Renal Tissue

**DOI:** 10.1101/632596

**Authors:** Brooke E. Harcourt, Aaron D. McClelland, Hiroshi Yamamoto, Hideto Yonekura, Yasuhiko Yamamoto, Sally A Penfold, Amelia K. Fotheringham, David A. Vesey, David W. Johnson, Melinda T. Coughlan, Mark E. Cooper, Phillip Kantharidis, Josephine M. Forbes

## Abstract

The *AGER* gene encodes for a number of RAGE isoforms, with the membrane bound signal transduction and “decoy” circulating soluble RAGE being the best characterised. Here we demonstrate a novel nuclear isoform of RAGE in mice and human kidney cortex which by cell and size fractionation we determined to be approximately 37kda. This nuclear RAGE isoform is functional and binds to DNA sequences within the upstream 5’ promoter region of its own gene, *AGER*. This binding was shown to be abrogated by mutating the DNA consensus binding sequences during electromobility shift assay (EMSA) and was independent of NF-□B or AP-1 binding. Cotransfection of expression constructs encoding various RAGE isoforms along with *AGER* gene promoter reporter-plasmids identified that the most likely source of the nuclear isoform of RAGE was a cleavage product of the nt-RAGE isoform. In obese mice with impaired kidney function, there was increased binding of nuclear RAGE within the A. Region of *ager* gene promoter with corresponding increases in membrane bound RAGE in renal cells. These findings were reproduced *in vitro* using proximal tubule cells. Hence, we postulate that RAGE expression is in part, self-regulated by the binding of a nuclear RAGE isoform to the promoter of the *AGER* gene (encoding RAGE) in the kidney. We also suggest that this RAGE self-regulation is altered under pathological conditions and this may have implications for chronic kidney disease.

## Introduction

The receptor for advanced glycation end products (RAGE) is a pattern recognition, multi-isomeric member of the immunoglobulin superfamily. RAGE is widely expressed and contributes to inflammation, diabetes and its complications, cancer, Alzheimer’s disease and a number of immunological conditions [1]. The RAGE gene (*AGER/ager*) is located in the Class III region of the major histocompatibility complex and encodes for endogenous secretory and membrane bound isoforms of RAGE, as well as numerous other tissue specific splice variants yet to be fully characterised [2–4]. In particular, the kidney is thought to express a high number of RAGE splice variants when compared with other organs, of which approximately 50% are thought to be targeted for degradation, but the function of the remaining splice variants remains unknown [3, 5]. Full length RAGE (fl-RAGE) represents a membrane spanning isoform which binds complex high molecular weight ligands including advanced glycation end products (AGEs), s100 calgranulins, beta amyloid and DNA bound high mobility group box protein-1 [6]. Ligation with fl-RAGE activates intracellular signalling pathways such as nuclear factor kappa B (NF-□B), [7] and JNK and JAK/Stat pathways, directly impacting on the transcription of many inflammatory genes.

Chronic increases in RAGE signalling in transgenic mice result in renal pathology [8], particularly in the context of stressors such as diabetes [9]. Blockade of RAGE expression or signalling has also been shown to prevent nephropathy in experimental models of diabetes [10–13], non-diabetic kidney disease [14–16] and obesity [17]. Not surprisingly, administration of sRAGE, a competitive decoy of RAGE is renoprotective in experimental models of chronic kidney disease. Circulating levels of sRAGE but not es-RAGE are predictors of albuminuria in diabetes [18], and are increased with obesity [19, 20].

Previous work from Tanji *et al*, performed using human renal biopsies presented photomicrographs that suggested the presence of nuclear RAGE [15]. Hence, the aim of the present study was to characterise this nuclear isoform of RAGE in the kidney and determine its relevance to renal pathology.

## Results

### RAGE deficient mice are protected from high fat induced renal dysfunction

Here it is demonstrated that RAGE deficiency afforded protection against the development of high fat induced renal injury. The rate of urinary albumin excretion was significantly increased with high fat feeding in wild type mice, but not in RAGE deficient mice (Figure 1A: AER). Similarly, creatinine clearance was increased in wild-type mice that were fed high fat diets indicative of renal hyperfiltration, and this was ameliorated in mice with RAGE deficiency (Figure 1B: CrCl).

**Figure 1:**
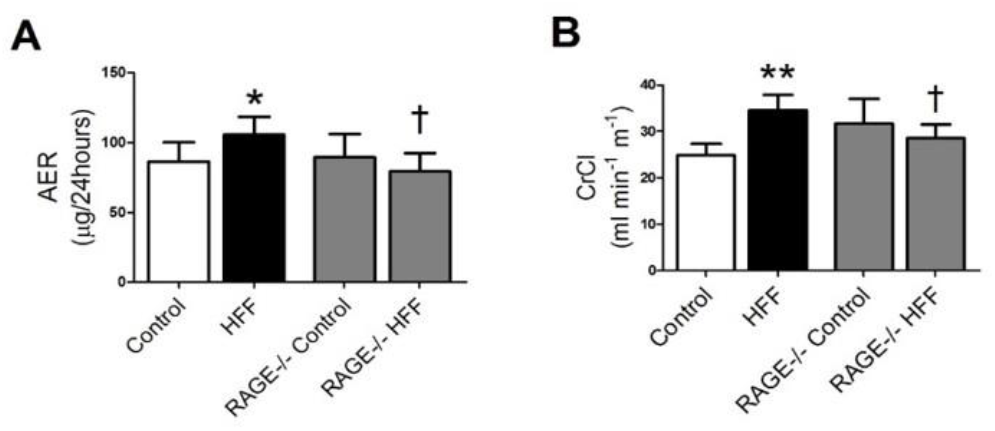
Renal Function in WT and RAGE Deficient (RAGE-/-) Mice. **A:** Urinary Albumin Excretion Rate (AER μg/24hours) by ELISA **B:** Creatinine clearance (CrCI ml/min/m^2^) by HPLC. High Fat Fed (HFF). \**p*<0.05 vs Control, \*\**p*<0.01 vs Control, ^†^p<0.05 vs HFF

### Nuclear RAGE expression is present in renal cortices

Membrane bound RAGE expression (flRAGE) was increased within renal cortices taken from obese mice with renal impairment (Figure 2A). For the first time, the presence of RAGE was also demonstrated within cell nuclei, from renal cortices of all wild type (WT) mice (Pictured Figure 2B-D), and this was further increased in obese high fat fed WT mice.

**Figure 2:**
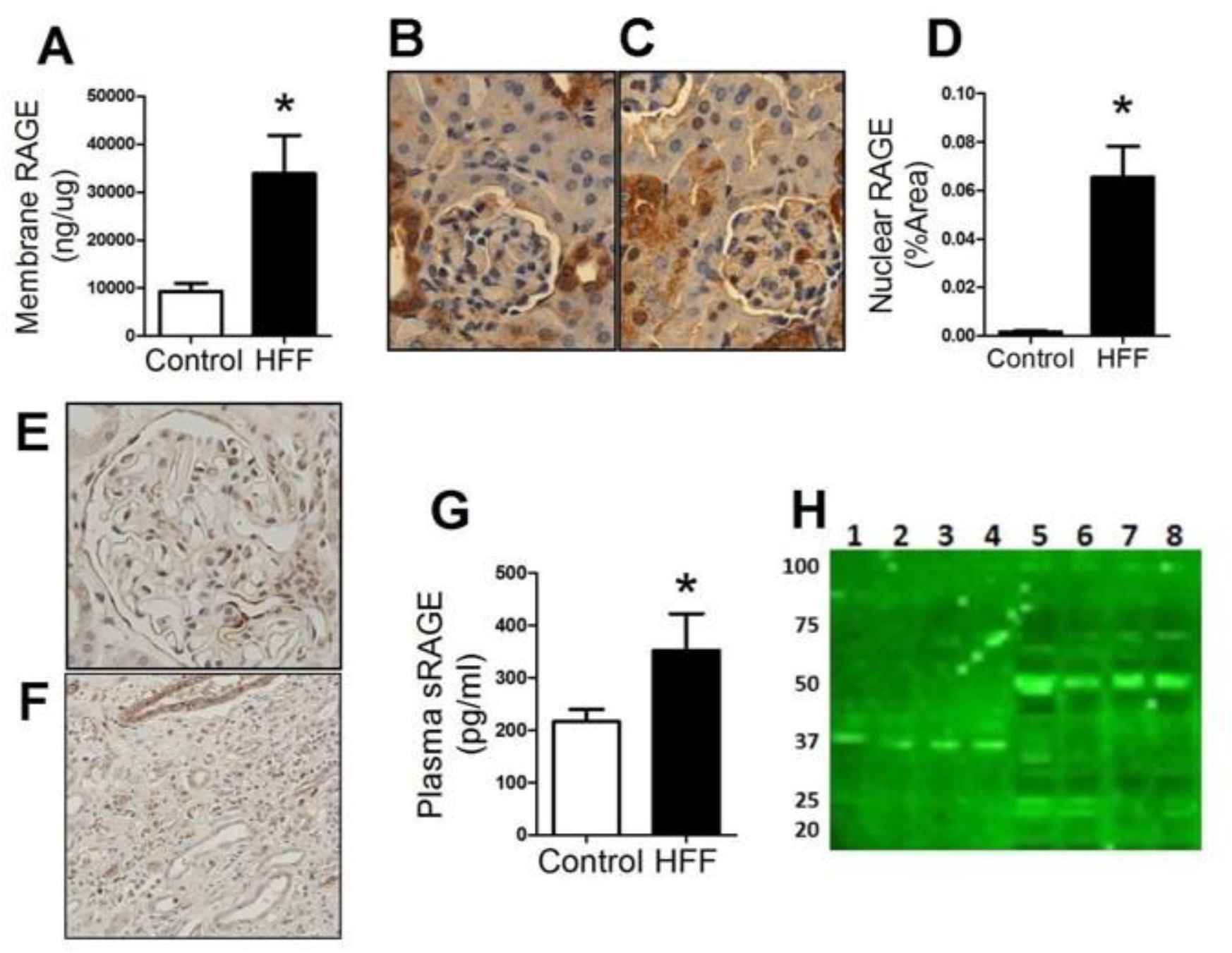
Expression of RAGE Isoforms. **A:** RAGE expression in membrane isolates from renal cortices of wild type Control and wild type HFF groups, as measured by ELISA. Representative immunohistochemical micrographs for **B.** WT Control mice, and **C:** WT HFF mice **D:** Quantification of RAGE positive stained nuclei as a percentage of total area, from WT Control and WT HFF mice. **E:** Representative micrographs for RAGE immunoactivity in Non-diabetic human renal nephrectomy. **F:** A human renal biopsy sample taken from an obese individual with T2DM. **G:** Quantification of soluble RAGE isoforms (sRAGE and esRAGE) in plasma from WT Control and HFF mice. **H:** Western immunoblot analysis of renal cortical fractions from WT mice, **NUCLEAR (LANE 1-4)** and **MEMBRANE (LANE 5-8).**High Fat Fed (HFF). \**p*<0.05 Control vs HFF

The presence of nuclear RAGE was confirmed in human nephrectomy tissues obtained from a healthy donor (Figure 2E), as well as within the renal cortex from an overweight individual with type 2 diabetes and nephropathy (Figure 2F).

Endogenous secretory RAGE (es-RAGE), which is also transcribed from ager/*AGER*, and is part of the circulating soluble RAGE pool (sRAGE), where the remaining sRAGE is contributed from membrane cleavage from endothelial cell surfaces [21]. With obesity, circulating soluble RAGE concentrations were increased as compared with control mice (Figure 2G).

Immunoblotting for RAGE in ultrafractionated nuclear extracts prepared from renal cortices of control and obese mice, identified a single 37kDa sized protein isoform, that was not identified in the other cellular fractions (Figure 2H: Lane 1-4). Immunoblotting for RAGE in the membrane fractions of kidney cortices from these mice showed full length (50kDa) and endogenous secretory RAGE immunoreactivity (25kda, Figure 2H), consistent with previous studies.

### Expression of Nuclear RAGE is inducible

Given the observation of RAGE protein within the nucleus, the capacity for RAGE to bind DNA was next examined using reporter constructs that contained known sequences of the *AGER* 5’promoter region to determine whether self-regulation of RAGE was occurring [22]. Indeed, RAGE has been reported to bind double stranded DNA in vitro (dsDNA, [23]) and CpG oligonucleotides in systemic lupus nephritis [24], although the reason for this is unknown. In the present study, rat proximal tubular cells were transiently transfected with a variety of human expression constructs to overexpress the known RAGE protein isoforms, es-RAGE, flRAGE, as well as N-truncated RAGE (ntRAGE). These were co-transfected with three different luciferase reporter constructs containing various known DNA binding elements within the *AGER* 5’ promoter region ([22]; Model Figure 3A).

**Figure 3:**
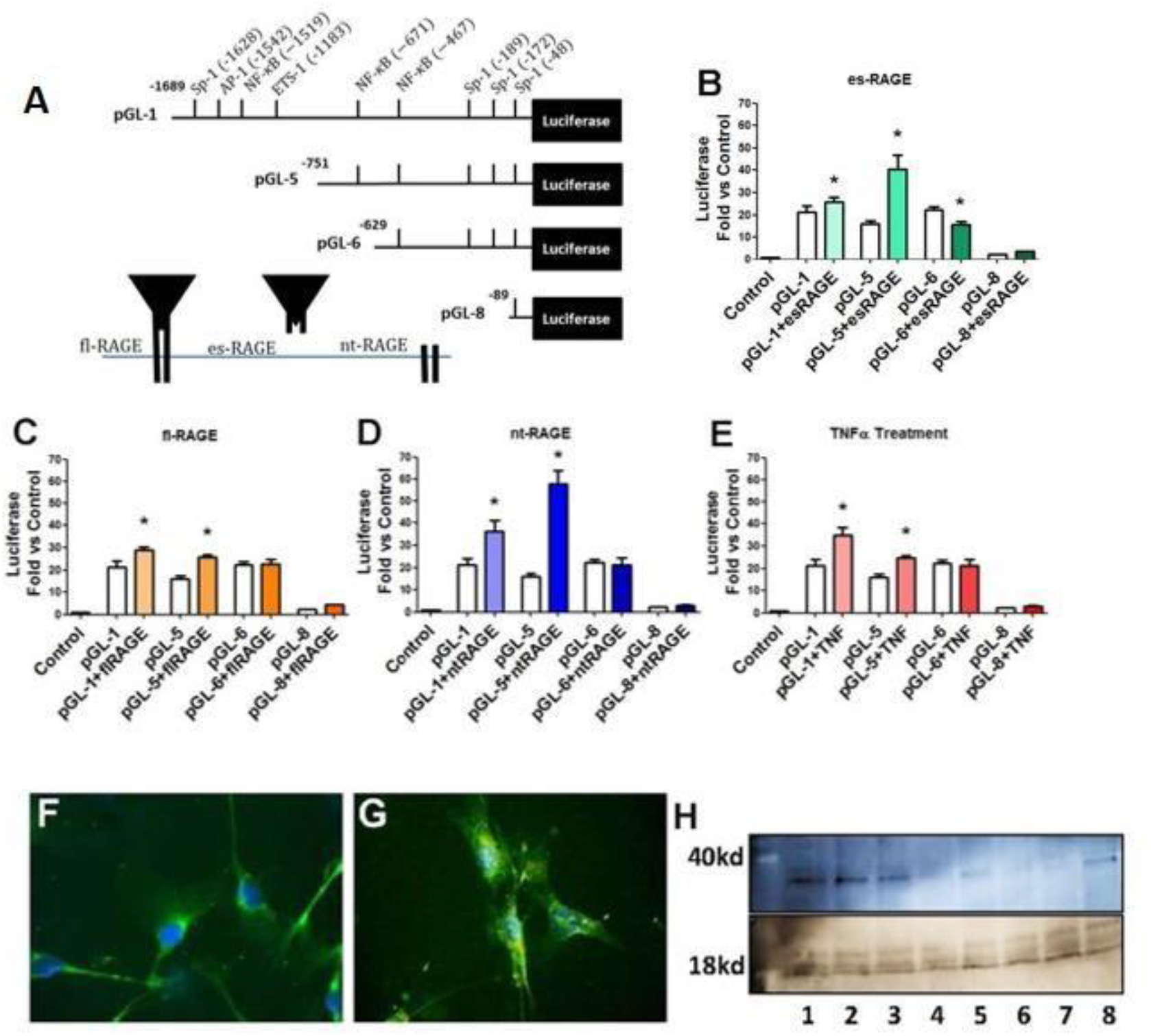
Inducible Expression of Nuclear RAGE. **A:** Model of luciferase reporter plasmids transfected into renal cell line containing various regions of the *AGER* gene (which encodes RAGE), to test transcription. **b-d** These reporter plasmids (PGL-1, PGL-5, PGL-6 or PGL-8) were cotransfected with plasmids containing cDNA to overexpress various RAGE isoforms.: **B**: Luciferase reporter plasmids for the 5’ promoter region of *AGER* co-transfected with es-RAGE over-expressing plasmid. Results represented as fold-change from Control. C: Luciferase reporter plasmids for the 5’ promoter region of *AGER* co-transfected with fl-RAGE over-expressing plasmid. Results represented as fold-change from Control. D: Luciferase reporter plasmids for the 5’ promoter region of *AGER* co-transfected with nt-RAGE over-expressing plasmid. Results represented as fold-change from Control. **E**: Luciferase reporter plasmids for the 5’ promoter region of *AGER* and treated with the *AGER* gene transcriptional activator TNF-α. Results represented as fold-change from Control. **f-g** Confocal micrographs of RAGE immunofluorescence (green) (using an antibody against the C-terminus of RAGE) nuclear staining (blue), in primary renal mesangial cells treated with **F**. Unmodified albumin or G. AGE modified albumin (AGE-BSA where nuclear staining is evident (green)), H: Western immunoblot of nuclear fraction of RAGE (~37kDa) from cells transfected with RAGE isoform over-expressing plasmids and histone-3 (16kda): LANE1-2 nt-RAGE, LANE 3-4 es-RAGE, LANE 5¬6 fl-RAGE, LANE 7-8 Control plasmid. \**p*<0.05 single transfection of plasmid vs double transfection of respective plasmid (white bar vs colour bar)

Firstly, transient overexpression of either fl-RAGE or nt-RAGE, co-transfected with the pGL-1 reporter, demonstrated an increase in *AGER* transcription, which was also seen following exposure to an inducer of RAGE transcription, TNF-α [22]. This region contains binding sites for Sp-1, AP-1 and NF-□Band activation of these transcription factors has been shown following RAGE ligation [25–27]. There was also robust induction of *AGER* transcription reported by the pGL-5 construct which contains NF-□Bbinding elements (nucleotides −751-0), following cotransfection with any RAGE overexpressing plasmid (Figure 3B-D).

This suggested that all of these RAGE isoforms have the capacity to bind within this *AGER* promoter region and induce RAGE transcription. It was previously shown that exposure of endothelial cells to AGE-BSA also led to DNA binding within the *AGER* promoter in regions upstream of that contained in the pGL-6 construct [22]. This suggested that RAGE ligands, such as AGE modified albumin, may potentially stimulate nt-RAGE or fl-RAGE production and translocation to the nucleus, as was seen in mesangial cells in the present study in response to AGE-albumin (Figure 3F and G). There was however, no change in reporter activity seen with either the pGL-6 (−629-0) or pGL-8 (−89-0) reporter constructs of the *AGER* promoter when cotransfected with plasmids overexpressing fl-RAGE or nt-RAGE. These data suggest that the fl and nt-RAGE derived isoforms may not bind within this region of the *AGER* promoter (−629-0). However, transient transfection to over-express esRAGE when co-transfected with the pGL-6 reporter construct of the 5’ *AGER* promoter (−629-0; Figure 3B), significantly decreased transcription, in contrast to that seen with both fl-RAGE (Figure 3C) and nt-RAGE (Figure 3D).

Furthermore, using western immunoblotting, it appeared that overexpression of nt-RAGE (Lanes 1-3; Figure 3H) resulted in elevated nuclear RAGE (Lanes 1-2) when compared with either fl-RAGE (Lanes 5-6. Figure 3H) or esRAGE (Lanes 3-4; Figure 3H).

### Chromatin interaction of nuclear RAGE in the 5’ promoter region of AGER

Given that other transcription factors have previously been identified following the binding of ligands to membranous fl-RAGE [25–27], the association of these factors with the binding of the nuclear RAGE isoform to DNA was examined. Using chromatin immunoprecipitation (ChIP) we identified that RAGE was bound to chromatin taken from nuclei of renal cortices of both control and obese mice (Figure 4A). RAGE bound to chromatin was increased in the A. Region (nucleotides 1628 to −1519) of the *ager* 5’ promoter region within renal tissue from obese mice (Figure 4A). This region also contained Sp1, AP-1 and NF-□B transcriptional binding elements. Whilst, RAGE bound chromatin within both the N. and S. Regions (Figure 4A-D) of the *ager* promoter, this was not different among WT and obese mouse groups. Nuclear translocation and DNA binding of Rel A (Figure 4E) and Rel B (Figure 4F) complexes of NF-□B, were also increased in kidney cortices with obesity, and were reduced in the kidneys of RAGE deficient obese mice (RAGE-/-).

**Figure 4:**
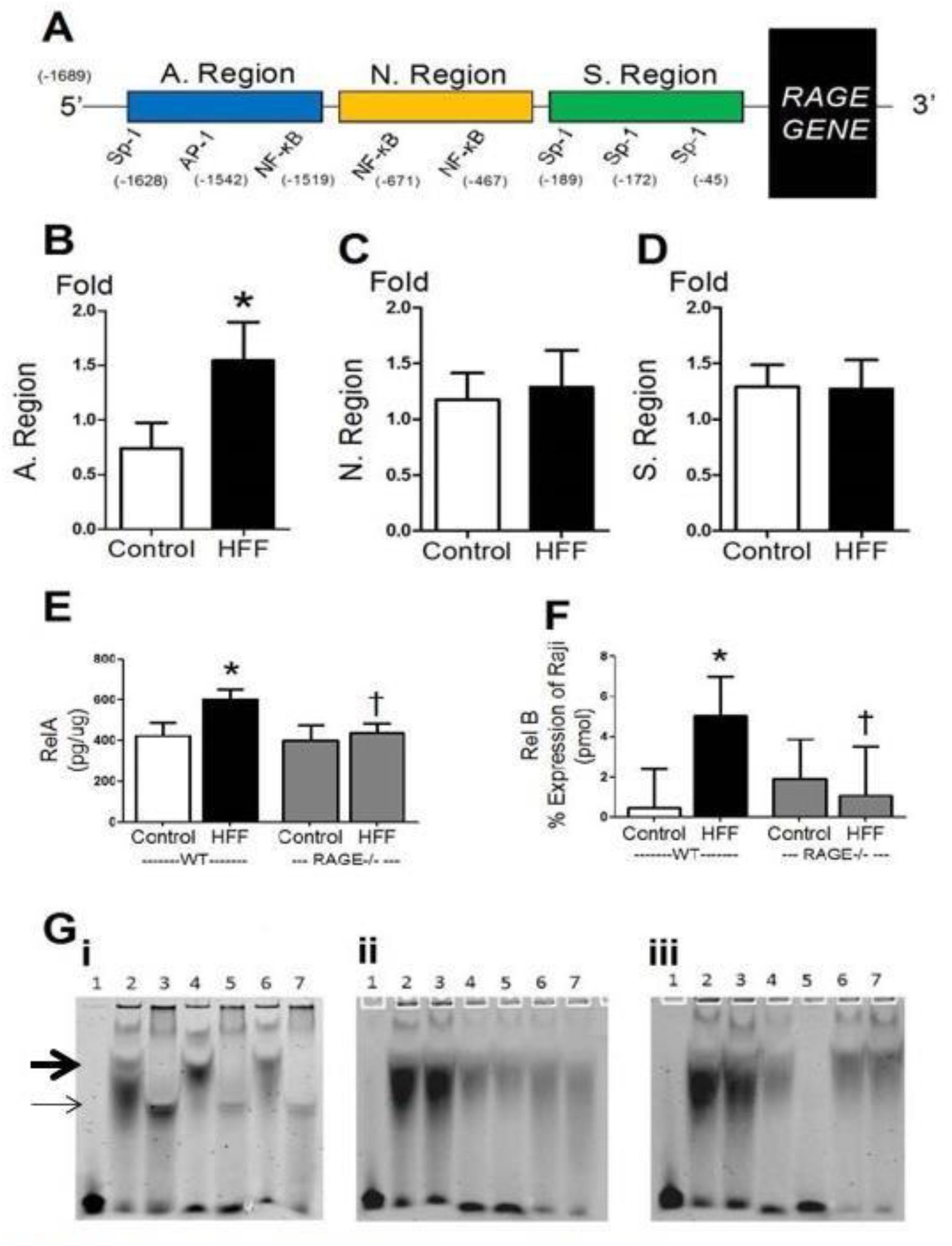
*AGER* (RAGE gene) Promoter Region Interaction and Transcription. **A:** Model of chromatin immunoprecipitation (ChIP) quantitative RT-PCR probe and primer design regions. A. Region (−1628 to−1519), N. Region (−671 to−467) and S. Region (−189 to −45) **b-d** Chromatin immunoprecipitation of RAGE on DNA from kidney cortices of WT Control and HFF mice. **B:** Interaction of RAGE with chromatin in the A. Region of the 5’ promoter region of ager. C: Interaction of RAGE with chromatin in the N. Region of the 5’ promoter region of **ager. D:** Interaction of RAGE with chromatin in the S. Region of the 5’ promoter region of RAGE. E: Total RelA DNA binding activity in renal cortices from WT and RAGE deficient mice. **F: Total** RelB DNA activity in renal cortex from WT and RAGE deficient mice. G: Electromobility shift assay (EMSA) of specific oliogonucleotide regions of the A. Region of the **ager** promoter in kidney cortex DNA from control mice demonstrating i) RAGE antibody Supershift, ii) Sp-1 antibody Supershift, iii) NF-κB antibody Supershift. **Lane 1.** No DNA Control. **Lane 2.** Construct 1 **Lane 3.** Construct 1. with Antibody Supershift, **Lane 4.** Construct 2. Lane 5. Construct 2 with Antibody Supershift, **Lane 6.** Construct **3, Lane 7.** Construct **3** with Antibody Supershift. H: Interaction of ER-ß bound chromatin in N. Region of the 5’ promoter region of the RAGE gene. I: Interaction of ER-ß bound chromatin in S. Region of the 5’ promoter region of the RAGE gene. High Fat Fed (HFF). RAGE deficient (RAGE-/-) \**p*<0.05 Control vs HFF, ^†^*p*<0.05 WT HFF vs RAGE-/-HFF

Using oligonucleotide sequences spanning the A. Region of the *ager* promoter in electromobility shift assays (EMSA) where increased chromatin binding of RAGE was detected during the ChIP analysis, protein interaction was further analysed. Increased DNA binding activity when the nucleotide sequence CACCCC with supershifting was confirmed using RAGE antibody (Figure 4Gi). This was confirmed using mutant oligonucleotide sequences (Suppl Figures 1A). Absence of DNA binding or antibody supershift was observed for Sp-1 and NF B (Figure 4Gii and iii).

Previous studies [22, 28], have also described the interaction of nuclear estrogen receptor ER-α with the *AGER* promoter region in human endothelial cells. Hence, the binding of ER-α and ER-β to chromatin was next investigated using ChIP, in renal cortices of WT and RAGE-/-mice with and without obesity. There were no differences seen between groups when ChIP for ER-α was examined within the *ager* promoter (Suppl Figures 1B-C), despite DNA binding of ER-α in all mouse groups. However, using ChIP, nuclei from renal cortices showed that estrogen receptor-β (ER-β) DNA binding activity within both the N. and S. regions of the *ager* promoter increased with obesity as compared with WT mice. This was not observed in RAGE-/-obese mice (Figure 4H-I).

## Discussion

For the first time, this study has demonstrated a novel nuclear isoform of RAGE in renal cortex and identified binding sites for nuclear translocated within the ager gene. The expression of nuclear RAGE was inducible by known ligands *in* vitro, and differentially by RAGE isoforms and was altered in a murine model of high fat feeding. The nuclear RAGE isoform was additionally identified in human nephrectomies.

The presence of RAGE in the nuclei of renal cortical cells is novel finding. Previously, this particular isoform may have been overlooked to the use of crude whole cell or tissue homogenisation and protein lysate analysis. However, some previous studies have suggested the presence of an additional functional RAGE isoform, with unknown localisation and function. Hudson *et al*, reported a ~40kDa RAGE protein isoform [2], which was produced as a result of genetic recombination of *ager*, but the cellular location of that isoform was not determined in that study. A RAGE isoform of ~40kDa was also described in total cell lysates of neuroblastoma cells [29], as well as peripheral blood mononuclear cells [30] but again the function and subcellular localisation of this isoform has remained undetermined. A similar sized functional protein was also identified by Kalea *et al* [3], in mice as a result of genetic recombination of *AGER*. In these studies the newly identified isoform of RAGE was not fully characterised for cellular location of function. These findings along with our present observations in the kidney suggest that further research into the function of this RAGE isoform and its effects on the transcription of other genes is warranted.

From the present study, it appears that already characterised isoforms of RAGE have varying effects on the transcription of the RAGE gene (ager). For example, overexpression of nt-RAGE, which is present of cell membranes but does not bind ARGE ligands, resulted in both increased RAGE transcription in reporter assay and increased expression of the nuclear RAGE isoform. Using specific oligonucleotide sequences in EMSA, the binding sites of nuclear RAGE in the AGER promoter were able to be characterised in renal tissue. In addition, supershift assay suggested that this nuclear RAGE was not acting by binding to known transcription factors to regulate *AGER* within this DNA region as distinct from other regions of the *ager* promoter. Indeed, there was no difference between control and high fat fed mice on ChIP for RAGE biding in regions other than the A region of the *ager* promoter. Other identified sites affecting *ager* transcription, were consistent with those previously identified for the *AGER* [31] Hence it is tempting to speculate that the nt-RAGE isoform, which in renal cells can be seen on the cell membrane surface but does not bind ligands and has an as yet uncharacterised role [4] is the major source of nuclear RAGE. Further, it is also suggested that this nuclear RAGE uniquely interacts with a binding site located within the A. Region of the ager promoter affecting the expression of RAGE in pathologies such as seen with obesity in this model. It was further identified that NF-□B and SP-1 remained unchanged in this region of the promoter.

Of interest, was the observation that cellular over-expression of other RAGE isoforms, such as membrane bound RAGE and es-RAGE, also caused transcription or suppression of ager gene transcription respectively. However, the over-expression of these particular isoforms did not appear to increase the appearance of RAGE in the nucleus which suggests that these RAGE isoforms likely caused effects on *AGER* via other transcription factors. Indeed, a number of previous studies, including our own [32, 33] have shown changes in gene transcriptional regulation via NF-κB and other transcription factors as was also seen in the present study for the NF-κB subunits, RelA and RelB. Interestingly, however *AGER* transcription induced by es-RAGE (nucleotides −629 to 0) was supressed in the present study. This is consistent with the thinking that soluble RAGE may regulate the expression or signal transduction of full length membranebound RAGE [34, 35], although the exact mechanism by which this occurred was beyond the scope of the present study. Taken together the findings highlight a complex mechanism of *AGER* gene regulation that has not been previously described.

The finding that RAGE can be found within nuclei and that it has direct binding activity within its own promoter lays foundations for the possibility that nuclear RAGE, most likely in association with proteins, such as NF-□B may play a regulatory role in the transcription of *AGER* and potentially other genes. There is also evidence provided within the present study that dysregulation of RAGE binding within its own promoter may occur during renal disease, contributing to pathology development and progression. These findings could directly translate to other diseases where RAGE is implicated as a pathogenic mediator.

## Methods

### Mouse Model

Male wild-type, C57BL/6J and RAGEdeficient (RAGE-/-, [14] backcrossed 9 times onto C57BL/6J) mice, were randomised to either control (AIN-93G, Specialty Feeds, Perth, Australia), or high fat diets (SF05-031, Specialty Feeds, Perth, Australia) diet for 16 weeks which resulted in significant weight gain (as previously described Control 2.9±2.4g vs high fat fed 11.0±1.6g, *p*<0.05, RAGE-/-Control 8.3±2.1g vs 15.4±2.2g, *p<*0.01[17]). Mice were monitored weekly for body weight and plasma glucose concentrations. Mice were housed in a temperate environment with a 12 hour light-dark cycle with *ad libitum* access to food and water. All studies were performed in accordance with the guidelines from the National Health and Medical Research Council of Australia and the AMREP Ethics Committee.

Renal function was determined using 24 hour urine and plasma collections at the study endpoint. Creatinine clearance (CrCl) and urinary albumin excretion rate (UAER) were determined, as previously described [11].

### Human Kidney Sections

Paraffin embedded renal sections were obtained from Clay Winterford (QIMR, Queensland, Australia). Normal renal tissue was obtained from patients undergoing partial nephrectomy for solid tumours.

### RAGE Protein Analysis

Kidneys were removed and immediately frozen in liquid nitrogen. The medulla was removed and the cortex was subjected to an ultrafractionation protocol that separated the cortex into membrane, cytosolic and the nuclear fractions, as previously described [11].

Cytosolic, membrane and nuclear protein fractions were analysed via ELISA (RAGE, Quantikine, R&D, Minneapolis, MN, USA).

The protein concentrations of the ELISA were corrected for the total protein concentrations of the sample as determined by Qubit fluorescence incorporation protocol (Qubit, Life Technologies, CA, USA).

Twenty-five micrograms of each renal cortex fraction was added to 10% or 12% polyacrylamide gels (Mini-protean TGX precast gels, Bio-rad, CA, USA), and assessed using infrared secondary probes (LICOR, NA, USA).

Immunohistochemistry analysis for RAGE protein was performed in paraffin embedded human and mouse kidney sections, as previously described [11].

Western immunoblotting was performed as previously described [11]. Briefly, 25ug of nuclear or membrane/cytosolic fractions from WT and WT high fat fed mice, were subjected to SDS-PAGE and western immunoblotting with rabbit anti-Cterminal RAGE (a kind gift from Prof M. Neeper, Novo Nordisk Industries).

Primary mouse mesangial cells grown on glass coverslips were exposed to AGE-BSA or BSA (100ug/mL [36], 72 hours) in normal growth media (DMEM, Invitrogen, CA, USA) Following fixation with cold methanol and blocking with 1% BSA, presence of RAGE in the nucleus was confirmed using immunofluorescence (primary antibody, 1:200 goat anti-RAGE (Millipore, CA, USA), 1:5000 secondary antibody donkey anti-goat 488 (LICOR, NA, USA)). Cells were mounted using fluroshield with DAPI mounting media (Sigma-Alderich, CA, USA) and imaged on an Olympus Xcellence Pro confocal microscope (20× magnification).

### Chromatin Immunoprecipitation (ChIP)

Kidney cortices were analysed for activity in the 5’ flanking region of the *ager* (RAGE) gene. Probes were designed using BLAST and were created to allow specific analysis of whether RAGE, ER-α or ER-β activity was able to directly influence the up-regulation of the RAGE gene. The primers designed also allowed for specific identification of the region of RAGE promoter that was bound. Promoter *A. Region* (−1628 to1519)*, Probe*; TCTGGAGATGTCAGCCC with an MGB quencher (Taqman, Invitrogen, USA), *Forward primer*; GTTCCCCACCCCACTTATATACTCT, *Reverse primer*; TCCCCATTTTTGGCATCTCT. Promoter N. *Region* (−671 to−467), *Probe*; CCCTCAGACACATCCTC with a 3’ MGB quencher (Taqman, Invitrogen), *Forward primer*; CAGCCCTGAACCCTTCATCTG, *Reverse primer*; CCCATGGTGACAGTCTTGAAGA. Promoter S. *Region* (−189 to −45*. Probe*; ACCTGAAGGACTCTTG with an MGB quencher (Taqman, Invitrogen), *Forward primer*; GGTCGGGTGAGATTGCTTCTAG, *Reverse primer*; TGCCAGGAATCTGTGCTTCTG.

Protein and DNA interactions in renal cortices were fixed in 2% paraformaldehyde, prior to chromatin extraction and purification. DNA fragments were sheared into 300bp fragments using Bioruptor sonication at 4°C (Ion Torrent, Life Technologies, NY, USA). Following preclearing with salmon sperm/Protein A agarose slurry (GE Healthcare, NY, USA), 1:12 for 1 hour at 25°C, chromatin was immunoprecipitated with ER-α (rabbit antiER-α, Millipore, CA, USA; 1:100), ER-β (rabbit anti-ER-β, Millipore, CA, USA; 4ug/ul), or RAGE (goat anti-RAGE, Millipore, CA, 1:50) at 4°C 18 hours with constant rotation. Protein bound DNA antibody interactions were removed with the additional salmon sperm/Protein A agarose. The bound, unbound, and input control DNA samples were purified and analysed. All results were expressed relative to values from WT Control, which were assigned an arbitrary value of 1. Statistical analysis was performed, and reported on Log-transformed data (Y=Log(Y)) as data was found to be nonparametric.

### Electromobility Shift Assay (EMSA)

EMSA was performed according to kit instructions (LICOR, NA, USA). Briefly, 10ug nuclear protein was incubated with dsDNA infrared tagged constructs (Table 1. 5’-3’ First strand, IDT, NSW, AUSTRALIA) with or without the additional supershift antiRAGE antibody (1ug, Millipore, CA, USA).

**Table 1:**
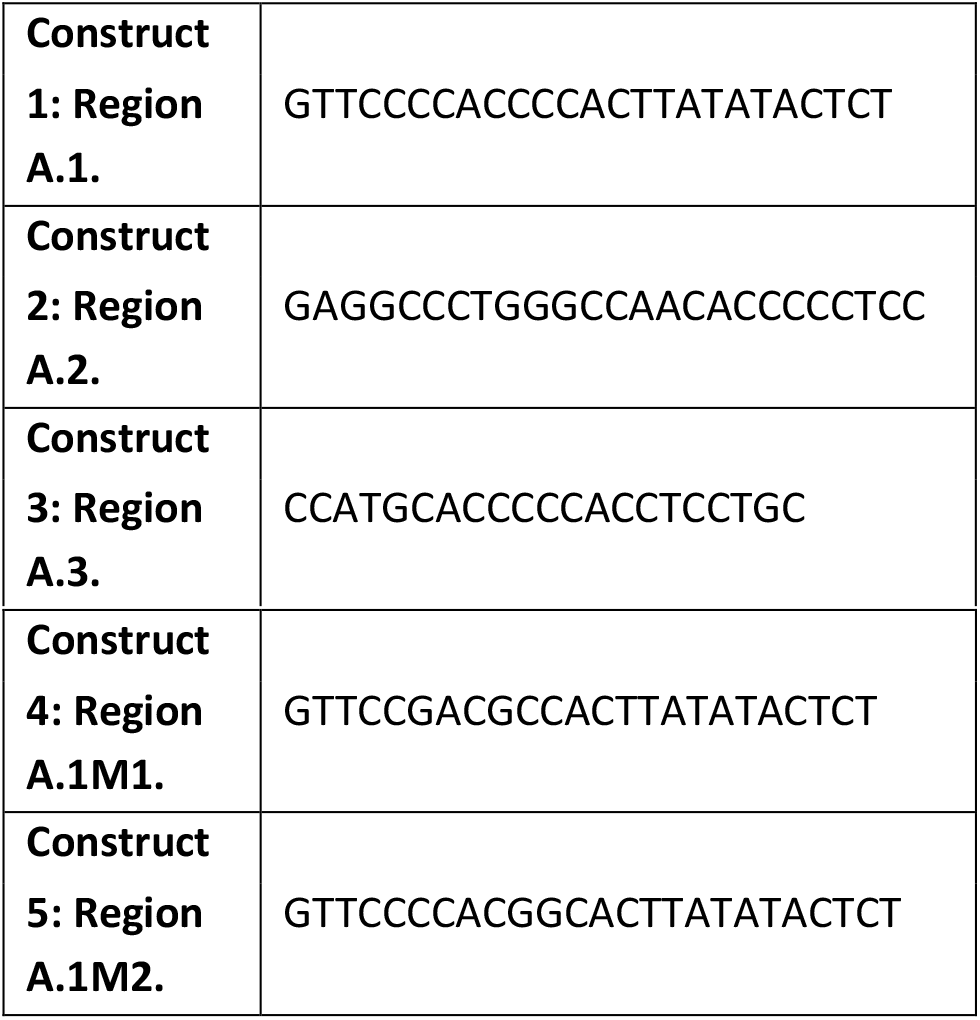
5’-3’ EMSA dsDNA constructs

### Luciferase Reporter Assay

Luciferase reporter assays for 5’regions of the *AGER* pGL-1, pGL-5, pGL-8, with cotransfection of cDNA plasmids to overexpress fl-RAGE, es-RAGE and nt-RAGE [22] were performed in the mesangial cell line; NRK-52E, in normal glucose conditions for 72 hours. Luciferase assays were performed as previously described [22].

### Statistical Analysis

Results are expressed as Mean (±SD) unless otherwise specified. Analyses were performed by ANOVA followed by, Tukey’s post-hoc analysis. Unpaired Student’s t-test analysis was used to compare means between 2 groups where indicated (Version 5, Graphpad Software, L Jolla, CA, USA). A *p*<0.05 was considered to be statistically significant.

## Acknowledgements and Declaration

The authors would like to acknowledge the technical assistance of Domenica Mc Carthy and Maryann Arnstein. We also thank Merlin C. Thomas and Mark A. Febbraio (Baker IDI Heart and Diabetes Research Institute, Melbourne, Victoria, Australia) for their assistance with the project. Further acknowledgement and thanks is paid to Professor Angelika Bierhaus (dec). Sections of paraffin embedded human kidney tissue were supplied by Clay Winterford (Queensland Institute of Medical Research. Brisbane, Queensland, Australia).

## Author Contributions

BEH, PK and JMF designed experiments, analysed data, wrote and revised the manuscript. ADM, HYa, HYo, YH were involved in the design and making of the constructs for the Luciferase Reporter Assays. SAP, AKF and DAM performed experiments. MEC and MTC edited the manuscript.

BEH is a NHMRC Peter Doherty Early Career Fellow, and JMF is an NHMRC Senior Research Fellow.

---

**Supplementary Figure.**
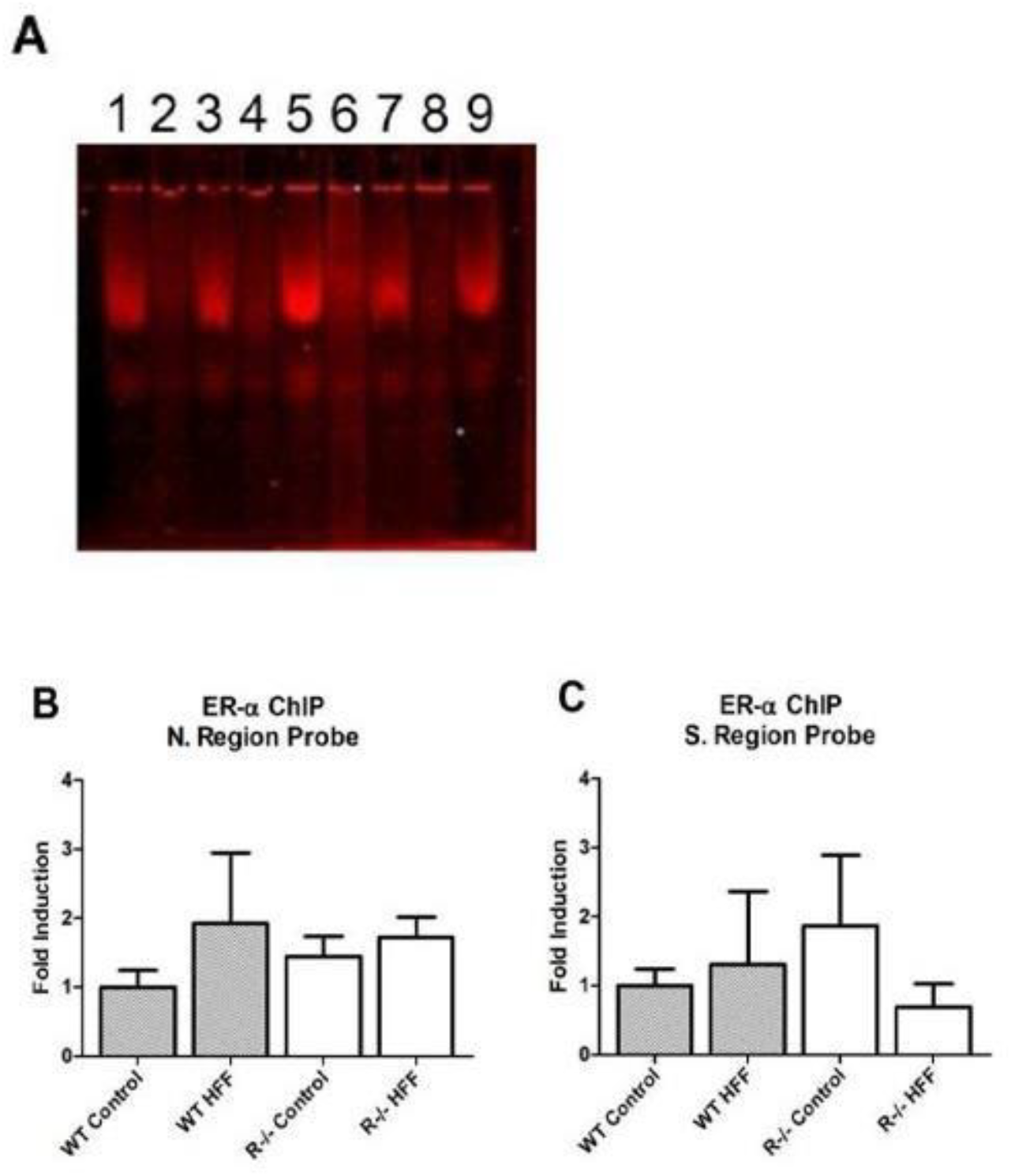
Electromobility shift assay (EMSA) of kidney cortex DNA from Control mice. **Lane 1 and 3.** Construct 4. **Lane 2 and 4.** Construct 4. with RAGE Antibody Supershift, **Lane 5 and 7.** Construct 5. **Lane 6 and 8.** Construct 5. with RAGE Antibody Supershift. **Lane 9.** No DNA, RAGE Antibody Control. **B-C** Chromatin immunoprecipitation (ChIP) of DNA from kidney cortex of WT and RAGE deficient. Control and high fat fed mice. B: Interaction of ER-a bound chromatin in A. Region of the 5’ promoter region of RAGE. C: Interaction of ER-a bound chromatin in N. Region of the 5’ promoter region of the RAGE gene.

